# Aptitude, not polyglotism, is associated with efficient activation in core language areas

**DOI:** 10.64898/2026.07.30.741766

**Authors:** Irene Balboni, Olga Kepinska, Alessandra Rampinini, Raphael Berthele, Narly Golestani

## Abstract

Understanding the cognitive architecture of the human language faculty requires exploring the boundaries of both predisposition and environmental experience. However, previous research on extraordinary multilingualism has often confounded language aptitude with multilingual experience, obscuring their distinct neural correlates. Here, we leveraged a linguistically diverse sample (N=121) and extensive behavioural testing to dissociate language aptitude from multilingual experience, modelling both dimensions continuously in whole-brain speech processing. Language aptitude and multilingual experience were weakly related, and their dissociation was also evident at the neural level. Higher language aptitude showed a neural signature of efficiency, characterised by lower activation in core perisylvian regions. In contrast, higher multilingualism was associated with greater engagement of regions implicated in narrative, multimodal, and memory processing, and with recruitment of traditional language hubs only during degraded speech processing, likely reflecting active attempts to decode unintelligible input. Finally, aptitude and experience interacted within sensorimotor regions. Continuous quantification of multilingual experience proved more sensitive than artificial grouping. By disentangling language aptitude from multilingual experience, this work provides a more precise account of the multilingual brain, and shows that its neurobiology can be better understood by modelling predisposition and experience as distinct but interacting dimensions.

## 1. Introduction

In neurobiological research, studying extreme linguistic talent provides a unique insight into the human language faculty, as exploring the boundaries of deficit and giftedness is fundamental to understanding its cognitive architecture. Researchers and laypeople alike have long been captivated by the mechanisms of language learning, the wide individual differences in language learning capacities, and the extreme levels of achievement possible within multilingualism. This is especially true when discussing’extraordinary’ multilinguals such as Emil Krebs, who reportedly spoke over 60 languages, or Cardinal Mezzofanti, who was credited with 51 ^1^. There is a near-universal fascination with polyglots that often stems from our own personal experiences and concerns about language learning, whether it is the struggle with our accent in a foreign language or the frustration of why some of us find second-language acquisition so difficult. However, neurobiological findings regarding polyglots, or what is considered’extraordinary’ multilingualism, are rare.

Early evidence came from a cytoarchitectural analysis of Emil Krebs’s brain. In a post mortem analysis of his brain, Amunts et al.^2^ identified significant differences in local connectivity and hemispheric asymmetries within Broca’s region compared to controls. More recently, fMRI research in the US has advanced this field by studying a combined cohort of 34 polyglots (defined as individuals who speak five or more languages, two of which they are highly proficient in) ^3,4^. Using a functional localiser designed to isolate linguistic syntactic and semantic processing, these studies found that polyglots exhibited reduced extent and lower magnitude of activation within perisylvian regions compared to controls, a finding interpreted as a signature of a “small and efficient language network”^3^ (p. 62), reflecting the optimisation of neural processing resulting from extreme linguistic experience. These findings are in line with the broader literature on talent and expertise, which suggest that expertise and intense practice lead to reduced and efficient activation within task-specific neural substrates ^5–7^. While these studies provide first insights into what are thought to be “unusual neurological resources”^1^ (p.28) of these very multilingual individuals, their conceptual and methodological choices may hinder our broader understanding of polyglotism and multilingualism in general.

Conceptually, previous studies of the polyglot brain have often relied on paradigms and methodological approaches to defining and operationalising polyglotism that may introduce systematic biases in the literature, with important scientific consequences. Much of the (neuro)linguistic research on polyglots has been conducted within WEIRD (Western, Educated, Industrialised, Rich, and Democratic) contexts, where monolingualism is often the institutional norm and linguistic minorities have historically been marginalised. This geographical and cultural context has dictated how we define *polyglotism*. Current literature portrays polyglots as “language super learners”^1^ (p. 5), setting them apart from other bilinguals or multilinguals due to the emotional and motivational factors that drive language learning ^8,9^ and their ability to speak at a level deemed exceptional in a largely monolingual setting. Typically, a cut-off point of five or six languages is used to define polyglotism ^1,3,8,10^. This creates two critical problems. First, these thresholds are geographically relative. While five languages may seem extraordinary in a largely monolingual WEIRD context, a repertoire of 3 or 5 languages or dialects is often a functional necessity for daily life in many multilingual regions ^11,12^. Second, because this level of multilingualism exceeds the’linguistic necessity’ of WEIRD communities, the participants recruited for these studies are, by definition, elective polyglots. These individuals are often expected to be gifted learners with a high aptitude for language^13^ (or simply aptitude, used interchangeably for brevity). This has traditionally been defined as an innate talent for language learning ^14,15^, particularly within formal settings ^16–18^. As a result, in current polyglot research, aptitude and multilingual experience might be confounded. While language learning aptitude is expected to predict outcomes for learners in formal settings, this relationship may not hold for circumstantial multilinguals^19^. In contexts where language acquisition is a functional necessity rather than a choice, aptitude likely plays a less central role than environmental demands.

More broadly, this limitation is related to the debate in cognitive neuroscience concerning the distinction between innate predisposition and experiential plasticity. The current literature on multilingualism and polyglotism is unable to determine whether rapid language acquisition is facilitated by a pre-existing biological blueprint (e.g. genetic profile), or whether this ability stems from learning and plasticity arising from years of intensive multilingual practice. Previous paradigms have frequently tested multilingual individuals who, due to the largely monolingual context, are likely elective multilinguals and therefore also likely talented learners. Consequently, it has been impossible to determine whether the polyglot brain is predominantly born or made. At the same time, most literature on bilingual brains relies on samples consisting mainly of circumstantial bilinguals. Participants with a migratory background or living in bilingual regions are tested, but no markers for talent are assessed. This creates a paradox in the current literature: although we know a great deal about circumstantial bilinguals and have some evidence from elective polyglots, we know much less about the neurobiology of circumstantial multilinguals and polyglots, as well as about neural markers of aptitude in less multilingual speakers. To answer these questions, we need a framework that considers language aptitude and experience as separate, potentially interacting factors rather than a single, combined trait. By conducting research in regions where multilingualism is the norm, we can sample both circumstantial multilinguals and elective polyglots, and potentially disentangle aptitude from multilingual experience at both behavioural and brain levels.

From a methodological perspective, previous neuroimaging studies of polyglotism have often relied on design choices that may inadvertently bias our understanding of both extreme multilingualism and language aptitude. First, previous studies have primarily relied on ROI-based analyses targeting cortical language hubs within the perisylvian region, such as Broca’s area^2^ and bilateral perisylvian regions^3,4^. However, a substantial body of work on second language learning and multilingualism has shown brain adaptations in regions implicated in executive control ^20–23^ and declarative memory ^24–26^. Furthermore, if multilingualism exerts broad effects on general cognition ^27–29^, ageing ^30–33^, and cognitive and neural reserve ^34^, these effects would be expected to extend beyond canonical language hubs. A whole-brain analysis is therefore needed to identify neural correlates of multilingualism in such domain-general regions. This is particularly important because, although multilingualism is the norm in many parts of the world, exceptional multilingualism, or polyglotism, may represent a cognitively demanding phenomenon that relies considerably on cognitive control and memory. Second, previous fMRI studies of polyglots have largely relied on subtraction-based contrasts, typically comparing intelligible speech or sentences with degraded speech or nonword stimuli, to isolate activation associated with linguistic processing. Although these contrasts are designed to capture phonological, lexical-semantic, and syntactic or combinatorial processes, they may also underestimate the extent to which nonwords ^35,36^ and degraded speech stimuli engage language-relevant processing. This is particularly problematic in polyglot and multilingual populations, as these individuals have access to broader linguistic repertoires, which may drive more effortful attempts to interpret degraded input. They may be more attentive to ambiguous speech-like stimuli and more motivated to extract meaning from them. Consequently, degraded input may engage components of the language network even when it contains no intelligible linguistic information per se. In line with this possibility, previous studies have reported significant IFG activation during degraded speech processing ^37^, particularly under conditions of active attention to the signal ^38^. Thus, if the control condition itself elicits meaningful language-related activation, the resulting contrast may effectively subtract out relevant linguistic responses, particularly in individuals with greater multilingual experience. This could contribute to the lower activation previously observed in polyglots in subtraction-based contrasts, such as speech versus degraded speech. Therefore, to accurately characterise the multilingual brain, it is necessary to examine responses to both intact and degraded stimuli, as well as the resulting contrast. Finally, previous studies of multilingualism and the brain have often relied on categorical group comparisons, imposing an artificial binary distinction between polyglots and controls that fails to capture the heterogeneous and graded nature of multilingual experience.

In the present work, we address these gaps by extending previous research on polyglotism to a more ecologically generalisable sample and a broader methodological framework. The study was conducted in Geneva, a highly multilingual context in which 42% of the population originates from outside Switzerland ^39^, and where heritage language use is therefore likely to be relatively common. Together with English and the two national languages studied in school curricula (German as the first foreign language and French as the local language of instruction), this context yielded a sample comprising both circumstantial (predominantly) and elective multilingualism. A substantial proportion of participants met the multilingualism threshold previously used to define polyglots, while the sample as a whole captured broad variation in language aptitude. We employed extensive behavioural testing to dissociate language aptitude from multilingual experience. This allowed us to sample broad variation in both aptitude and multilingualism, and to examine whether these two dimensions are associated with distinct neural correlates. Moving beyond binary group comparisons, we quantified multilingualism using a continuous measure and employed whole-brain analyses to identify neural correlates of multilingual experience both within and beyond traditional cortical perisylvian regions. Further, we examined neural responses not only for what is conventionally referred to as the high-level language contrast in the language localiser, that is, intact versus degraded speech stimuli, but also for each of the two conditions used to derive this contrast. This allowed us to test whether highly multilingual and/or high-aptitude individuals would use additional resources to process degraded, unintelligible input. We hypothesised that higher levels of multilingualism may be associated with reduced activation in traditional high-level language contrasts^3,4^, and that examining the constituent intact and degraded speech conditions separately would help clarify whether degraded speech elicits language network recruitment in more multilingual individuals. Finally, we predicted higher activation during intact speech processing and in the subtraction contrast measuring activation specific to phonological and high-level speech features within domain-general areas that have previously been shown to be modulated by multilingualism.

## 2. Results

### a. Multilingualism and aptitude: behavioural profiles

The sample exhibited a high degree of linguistic diversity, reflecting its unique demographic makeup. Figure 1 illustrates the wide variability in multilingualism across the sample, highlighting the continuous nature of these linguistic profiles. To test whether our primary predictors could be dissociated, we examined the relationship between language aptitude and multilingual experience using correlation and linear analyses. As shown in Figure 1, both multilingual experience and language aptitude were broadly distributed and only weakly correlated. Moreover, although the multiple regression analysis (Table 3) revealed that language aptitude significantly predicted multilingual entropy (β= 0.038, *p* =.022), the effect size indicates that these two constructs remain functionally dissociable. Specifically, the unique contribution of aptitude to the variance in multilingual experience was small, especially when compared to the much stronger effect of age (β = 0.050, *p* <.001). This weak association demonstrates that, in this highly multilingual context, language learning aptitude and multilingual experience are dissociable. For example, as shown in the scatterplot in Figure 1A, our sample included individuals with lower multilingual exposure who nonetheless possessed exceptional aptitude scores, and vice versa. Additional information on the descriptive statistics for the group derived from the binary categorisation can be found in the Supplementary Materials (Figure s1, Table s1).

**Figure 1.**
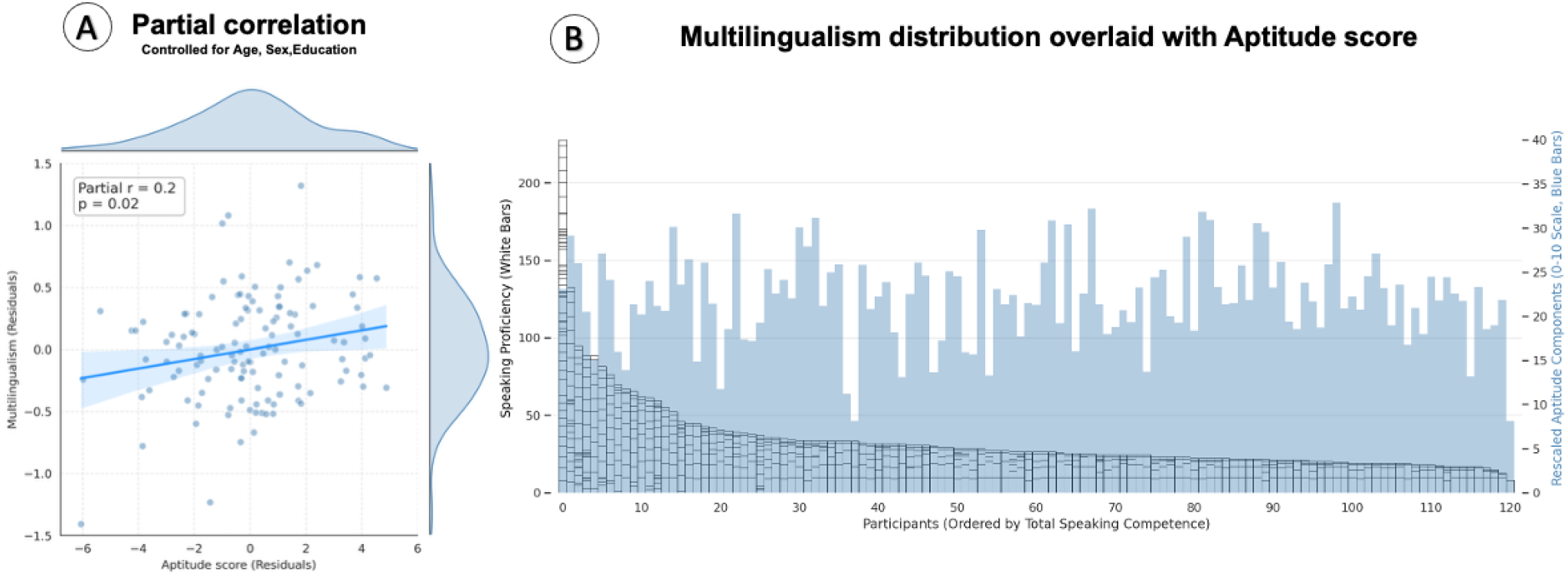
Distribution and relationships between language aptitude and multilingualism scores. Panel A: Partial correlation between language aptitude and multilingual experience (both z-scored), controlling for age, sex and education. Density plots represent the marginal distributions of each variable. Panel B: Multilingualism distribution (white stacked bars) overlaid with Aptitude scores (light blue). Multilingualism is represented by stacked segments within each bar, which indicate self-reported speaking proficiency in individual languages; thus, total bar height reflects cumulative speaking proficiency across languages, whereas the number of segments reflects the number of languages reported.

Two additional multilingual experience indices are provided: total number of languages known at any level and number of languages known at advanced proficiency, defined as total self-reported score ≥ 24/30 across reading, comprehension and speaking.

Multilingualism is operationalised as entropy score based on current speaking proficiency, and language aptitude as a compound score obtained from traditional aptitude measures (*N*=121)

### b. Neural correlates of multilingualism defined continuously and categorically

Figure 2 illustrates the whole-brain activation patterns for intact speech, degraded speech, and the high-level speech contrast across the entire sample. Significant results from the voxel-wise whole-brain regression analysis (full interaction model) across all functional contrasts are detailed in Table 4, which provides a comprehensive overview of the results from both the continuous (multilingual entropy and language aptitude) and categorical (group) analyses. The single-predictor analyses broadly converged with the full model (see Table s2 in the Supplementary Materials), while also revealing additional effects reported in the Supplementary Materials. Unless otherwise specified, the results discussed below refer to the full interaction model.

**Figure 2.**
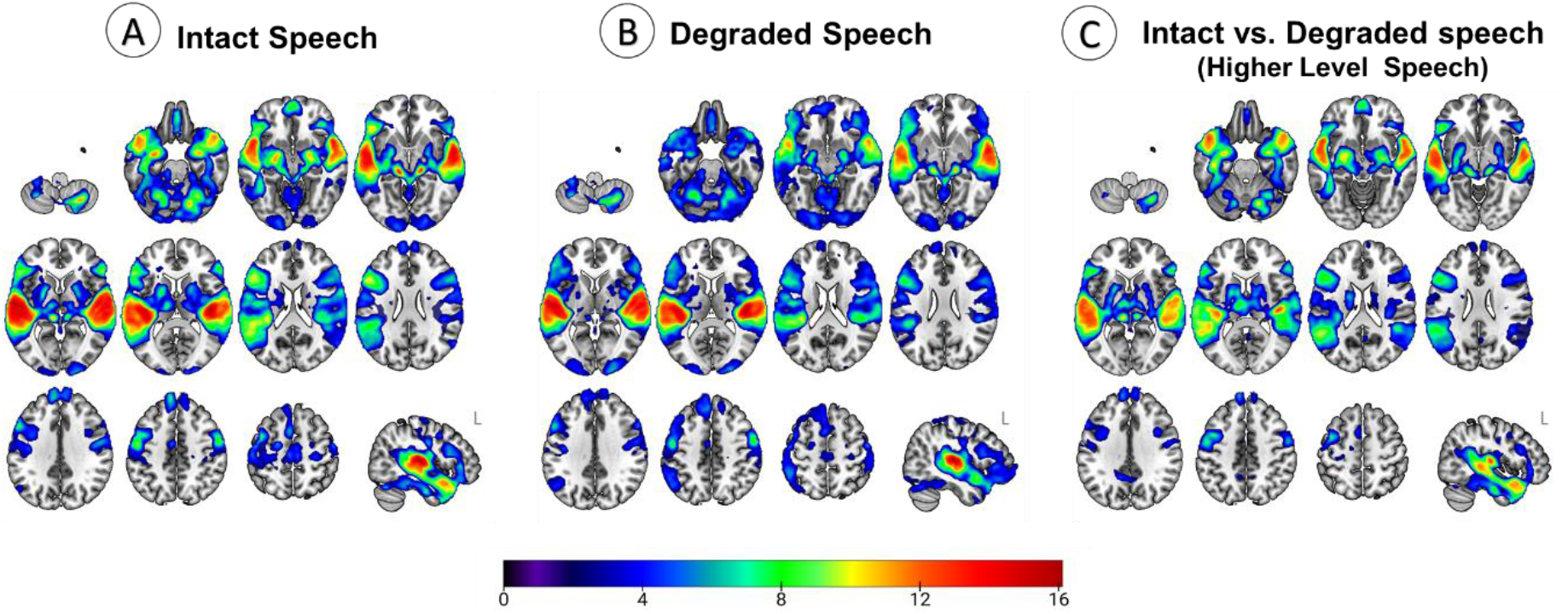
Whole-brain activation patterns for intact speech, degraded speech, and the high-level speech contrast across the entire sample. The colour bars indicate voxel-wise Z-score values for significant neural activation. Statistical images were thresholded using a cluster-forming threshold of Z > 2.3 and a cluster-corrected significance threshold of *p* =.05. L= left hemisphere.

The whole-brain categorical analysis comparing polyglots and controls revealed only a single significant effect: a cluster in Crus I of the left cerebellum, where controls exhibited greater activation than polyglots during the degraded speech condition (Figure s3). Although visual inspection suggested different activation levels across the three activation estimates (Figure s2), with higher activity in controls than in polyglots, these differences were not statistically significant. By contrast, when multilingualism was operationalised continuously using the entropy score, we observed widespread associations with neural activation across all three contrasts (Figure 3, Table 4). The patterns are described in detail in the next section.

**Figure 3.**
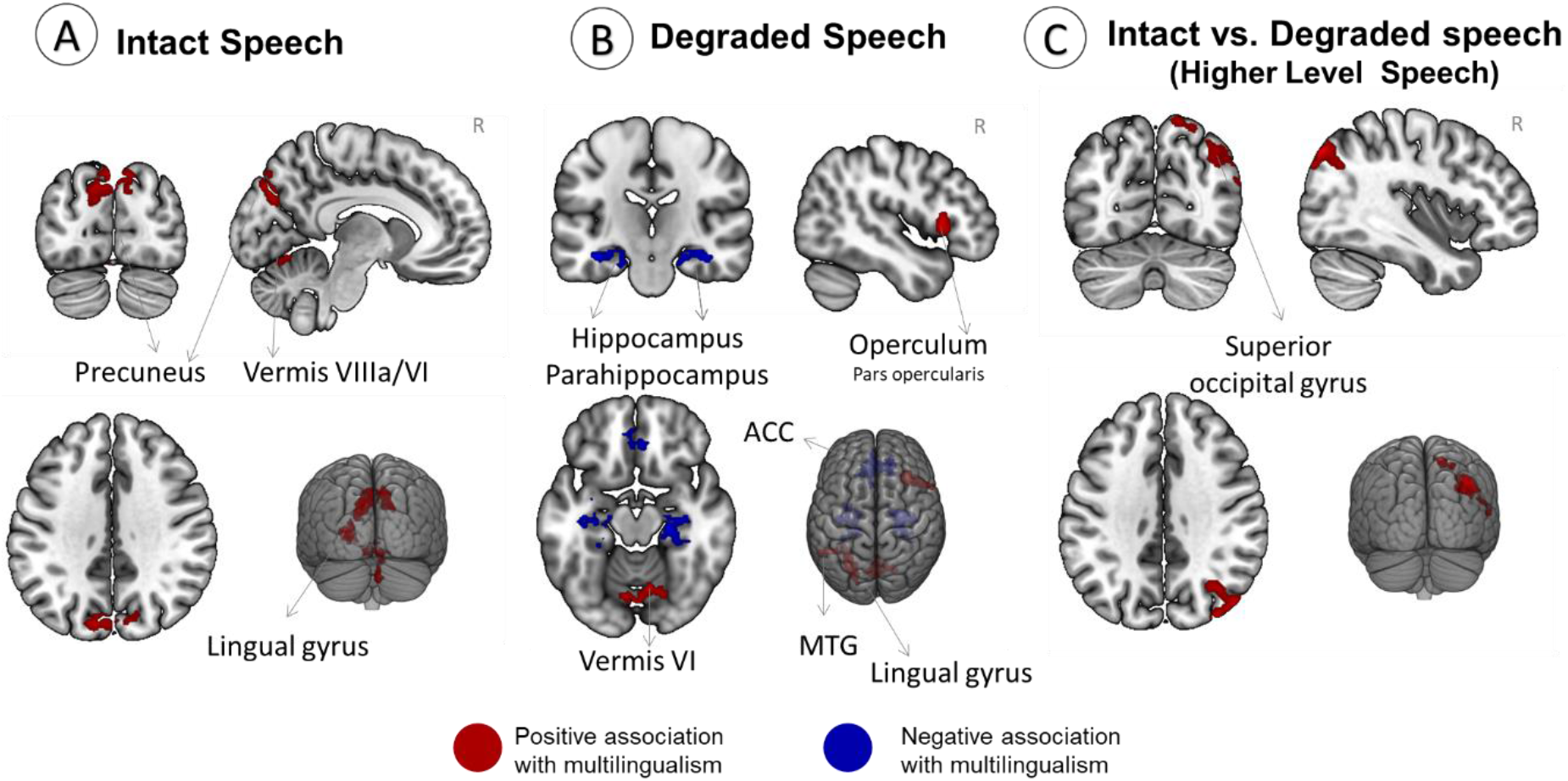
Whole-brain activation differences associated with continuous multilingualism scores. Red indicates positive associations with multilingualism, whereas blue indicates negative associations Statistical images were thresholded using a cluster-forming threshold of Z > 2.3 and a cluster-corrected significance threshold of *p* =.05. R= Right hemisphere

Furthermore, the categorical comparison in the group-constrained subject-specific (GSS) fROI analysis, performed to assess whether previous findings could be reproduced in our sample, yielded no significant differences for any contrast in any of the evaluated cortical language ROIs, with the exception of one uncorrected effect (p=0.045) in the right anterior superior temporal gyrus (STG) (full results in Table s3).

### c. Neural correlates of multilingualism

During intact speech listening (Figure 3A), higher multilingual entropy scores (henceforth: higher multilingualism) were only associated with increased activation in posterior and cerebellar regions, specifically the left precuneus and lingual gyrus, and the right cerebellar vermis VIIIa/VI.

No regions showed a negative association with multilingualism. In the high-level speech contrast (Figure 3C), higher multilingualism was associated with greater activation in right superior occipital/lateral parietal regions and, in the single-predictor models only, right hippocampal and parahippocampal regions.

In the degraded speech condition, higher multilingualism was associated with increased activation in several perisylvian and posterior regions, particularly the right IFG (pars opercularis), left middle temporal areas, and the lingual gyrus (Figure 3B). This effect was also observed in the left IFG in the single-predictor model (Table s2). Conversely, more multilingual speakers showed reduced activation in the anterior cingulate cortex (ACC) and bilateral hippocampal and parahippocampal regions during this condition.

### d. Neural correlates of language aptitude

During intact speech listening (Figure 4A), individuals with higher aptitude scores showed significantly lower activation in the left supramarginal gyrus (SMG), angular gyrus, and middle frontal gyrus (MFG), extending into the frontal pole. This negative association with aptitude was even more pronounced during the high-level speech contrast (intact > degraded; Figure 4C). High-aptitude participants exhibited markedly lower activation across a broad perisylvian and temporal network, including the left IFG, bilateral STG, planum temporale (PT), HG, and left planum polare (PP).

**Figure 4.**
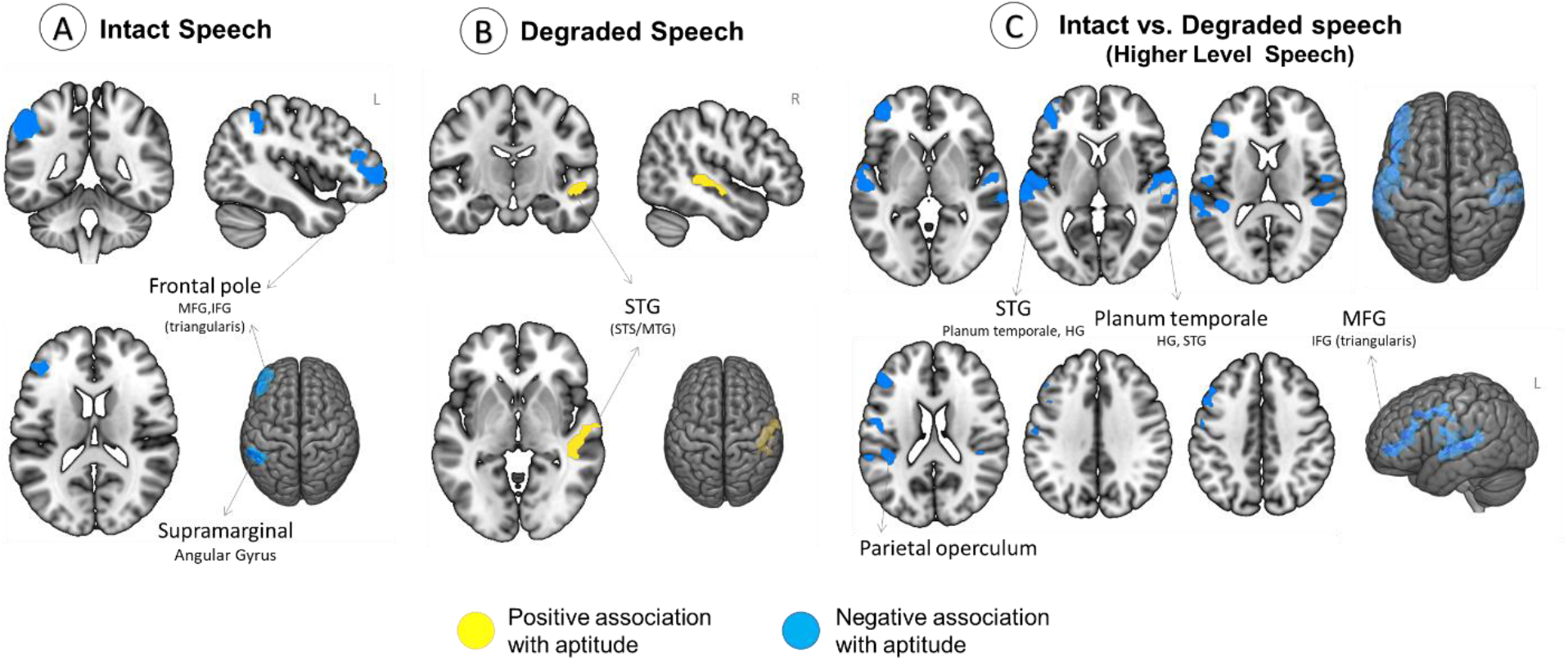
Whole-brain activation differences associated with aptitude scores. Yellow indicates positive associations with aptitude, whereas light-blue indicates negative associations. Statistical images were thresholded using a cluster-forming threshold of Z > 2.3 and a cluster-corrected significance threshold of *p* =.05. R= right hemisphere, L= left hemisphere

The degraded speech condition (Figure 4B) revealed positive associations between language aptitude and activation in the right middle and posterior STG and STS/MTG.

### e. Interaction between aptitude and multilingualism

To determine whether the neural correlates of multilingualism are modulated by individual differences in language aptitude, or vice versa, we evaluated the whole-brain interaction effect between multilingual entropy and language aptitude scores (Figure 5). Significant interaction effects emerged solely during the Intact Speech and Degraded Speech listening conditions. No significant clusters passed the threshold for the high-level speech contrast (Intact > Degraded).

**Figure 5.**
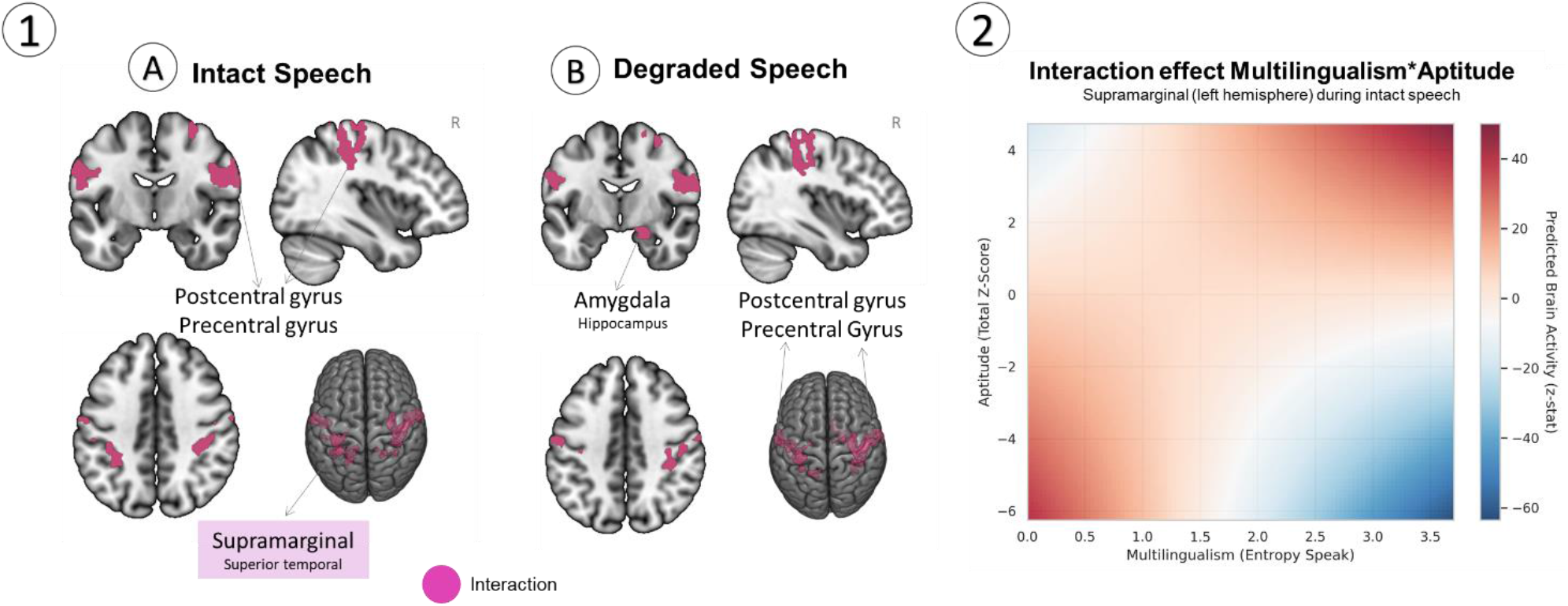
Interaction between multilingual experience and language aptitude in functional brain activation. (1) Whole-brain statistical parametric maps displaying significant clusters for the interaction between multilingual experience and language aptitude. Statistical images were thresholded using a cluster-forming threshold of Z > 2.3 and a cluster-corrected significance threshold of *p* =.05. Only conditions showing a significant interaction effect are displayed. R= right hemisphere. (2) Post-hoc decomposition of the interaction effect within a representative significant cluster (highlighted in panel A). Predicted neural activation within the significant ROI is plotted as a function of continuous multilingual entropy and aptitude score

During intact speech processing (Figure 5, Panel 1A), an interaction emerged in the right postcentral gyrus extending into the superior parietal lobule and the left SMG extending into the superior parietal lobule. Two additional clusters were isolated in the bilateral postcentral gyri. A similar, positive interaction pattern emerged when participants listened to acoustically degraded speech (Figure 5, Panel 1B). Moreover, the degraded speech condition uniquely engaged a subcortical limbic cluster spanning the right amygdala and part of anterior hippocampus.

Figure 5 (Panel B) shows the interaction between aptitude and multilingualism in a single representative cluster, as all significant interactions followed a similar pattern (plots for all significant clusters are shown in the Supplementary Materials Figure s4). The relationship between multilingual experience and neural activation depended on language aptitude, and vice versa. The interaction pattern suggests that neural activation was highest when aptitude and multilingual experience were concordant at the extremes of the sample, that is, in individuals with high levels of both or low levels of both, whereas discordant profiles showed lower activation.

## 3. Discussion

### a. Language aptitude and multilingualism are dissociable constructs with different brain correlates

The present study sought to clarify how multilingual experience and language aptitude, two constructs often intertwined in research on polyglotism, are related behaviourally and neurally. As argued in the introduction, disentangling the neural signature of language aptitude and multilingual experience is essential for fully understanding both phenomena. Previous studies on polyglots and bilinguals have often confounded these two factors. In our work, by leveraging a setting characterised by high ambient multilingualism, we were able to dissociate the neural correlates of these two dimensions across continuous variation in both constructs. Our previous behavioural analysis of this dataset already indicated a dissociation between language aptitude and multilingual experience in largely the same participants as those tested here. Using Exploratory Graph Analysis, we demonstrated that these constructs consistently dissociate into distinct behavioural clusters ^42^. In the current study, this divergence was further supported by the low correlation and limited shared variance observed between the two variables. This weak association does not preclude stronger relationships in other samples; indeed, previous work has reported a modest association between language aptitude and multilingual experience in participants selected for average-to-high aptitude and likely with more elective multilingual profiles ^48^. Nevertheless, in the present sample, the limited shared variance between aptitude and multilingual experience supports the view that these dimensions are fundamentally dissociable. Language aptitude likely reflects individual differences in predisposition for language learning and, while showing some modifiability through learning and ageing ^49,50^, can be considered a relatively stable trait ^16,51^, particularly in adults who may have learned languages through a mixture of formal instruction and immersion. Multilingualism, on the other hand, is a measure of experience and, even if language aptitude can certainly drive individuals toward linguistic “niche-picking”, that is, actively seeking out opportunities to learn and use new languages, environmental constraints often play a strong role in shaping linguistic profiles. In our sample, this resulted in significant variability in aptitude-multilingualism profiles. This included the less discussed high-aptitude individuals who predominantly use only one language due to a lack of environmental necessity or exposure, and low-aptitude individuals who achieve high levels of multilingual experience because of functional demands. The dissociation observed at the behavioural level is mirrored in our neuroimaging results, which reveal spatially distinct neural correlates for each construct. The neural correlates of aptitude are heavily anchored within core language cortical hubs, specifically left-lateralised frontal and parietal regions during intact speech, and frontal and temporal/auditory structures during the high-level speech contrast. In contrast, the effects of multilingualism were observed primarily outside the core language network for the two intelligible speech conditions.

Interaction effects were detected within the intact and degraded speech conditions, revealing highly overlapping clusters across conditions in the precentral and postcentral gyri. These interaction effects require cautious interpretation, given the complexity of the patterns obtained and because, to our knowledge, this is the first study to directly model the aptitude–multilingualism interaction in speech processing. This spatial convergence across conditions suggests that the interaction is likely to be driven by features common to both conditions, with exceptions localised to the left supramarginal cortex during intact speech and the right amygdala/anterior hippocampus during degraded speech, which might relate to condition-specific characteristics. The pre-and postcentral regions may therefore reflect responses to human-generated speech and speech-like stimuli. These regions have been shown to display bilateral selective activity for human-produced vocalisations over tones or natural sounds ^52^ and to be involved in speech-in-noise tracking ^53^. Furthermore, these sensorimotor networks contribute functionally to the perception of speech sounds ^54^ and are central components of a system coding for articulatory features such as voicing, place, and manner of articulation ^55^. Regarding the shape of the interaction, our findings suggest a concordance pattern, with participants showing matched profiles on our measures, high aptitude/high multilingualism or low aptitude/low multilingualism, exhibiting higher activation than those with discordant profiles. This pattern suggests that language aptitude modulates the relationship between multilingual experience and activation in sensorimotor speech-related cortices. In the post hoc visualisation, multilingual experience was positively associated with activation among high-aptitude participants, negatively associated with activation among low-aptitude participants, and only weakly related to activation among participants with average aptitude. This pattern suggests that the neural consequences of multilingual experience may depend on the aptitude profile in which that experience is embedded, rather than exerting a uniform effect across individuals. Within pre-and postcentral regions, which have been implicated in speech perception and articulatory-feature coding, this may reflect different routes to processing speech and speech-like auditory inputs: greater sensorimotor engagement in individuals combining high aptitude with high multilingual experience, but reduced recruitment with increasing multilingual experience among lower-aptitude individuals. Because this interpretation is based on post hoc visualisation of a continuous interaction, it should be treated as exploratory.

### b. Multilingualism modulates extra-perisylvian activation during intelligible speech and both perisylvian and extra-perisylvian activation during degraded speech

Our continuous analyses examining the neural correlates of multilingual experience revealed a functional pattern situated predominantly outside traditional perisylvian language regions. For the two conditions/contrasts tapping into intelligible speech processing, i.e. intact speech and high-level speech contrast, higher multilingual entropy scores consistently predicted increased neural recruitment. The notable exception, both in terms of the directionality of the effect and the specific regions involved, was the degraded speech condition, where multilingual experience was associated with both increased and decreased activation across different regions.

During intact speech processing and in the high-level speech contrast, higher multilingualism was associated with higher activation in extra-perisylvian regions, mostly in regions implicated in lexical processing, narrative tracking, and reading-related processes. In the intact speech condition, higher multilingualism predicted greater activation in the bilateral precuneus, an area important for lexical retrieval ^56^, lexical information processing ^57^, and verbal memory processes in bilinguals, particularly in the first language ^58^. Moreover, the precuneus is part of the dorsal regions of the default mode network ^59^ and is a key node for tracking longer narrative comprehension ^60^ and mental story representation ^61^, with its connectivity to language areas being predictive of language performance ^62^. Considering that our stimuli consisted of story passages (excepts from *Alice in Wonderland*), this higher activation may reflect greater engagement of story tracking and lexical processing mechanisms in more multilingual participants. We also observed higher activation in the left lingual gyrus, a visual area participating in letter and word decoding ^63,64^, although its functional activation is not exclusive to words ^65^. This area has been shown to be active in sentence comprehension across languages in bilinguals ^66^. Finally, higher activity was detected in more multilingual individuals in the right cerebellar vermis, a region connected to sensorimotor and attentional networks ^67^.

In the high-level speech contrast, the increased activation within superior occipital clusters in more multilingual individuals represents an unexpected result. A cautious interpretation could link this finding to evidence that these regions are involved in gesture and sign-language processing in hearing non-signers ^68^. More broadly, multilinguals are exposed to a broader multimodal repertoire, including language-and context-dependent use of gesture ^69^. Developmental work further suggests that bilingual exposure is associated with increased attention to mouth and face cues during infancy ^70,71^, while studies of multilingual classroom interaction indicate that gesture may provide an additional communicative resource at low proficiency levels ^72^. Together, these findings raise the possibility that the occipital/lateral parietal effect observed here reflects greater engagement of multimodal processing infrastructure in more multilingual individuals. This interpretation remains speculative, as the present task involved auditory speech only and did not directly manipulate gesture, face, or visual-speech cues.

Moreover, the single-predictor model revealed that multilingual individuals exhibited greater activation in the right hippocampal and parahippocampal regions during high-level speech processing. The hippocampus has long been recognised as a core structure for lexical learning and the declarative memory processes underlying vocabulary acquisition ^73–75^. Furthermore, it has been directly linked to neuroplasticity during intensive language training; for instance, a seminal longitudinal study by Mårtensson et al.^26^ on interpreters found that while bilateral hippocampi increased in volume after three months of intensive learning, only the right hippocampus showed more prominent changes in learners with higher foreign language proficiency outcomes, possibly indicating a combined effect of language learning and higher aptitude. In the present, more heterogeneous sample, the right hippocampal effect emerged in relation to multilingual experience in the single-predictor model, which is consistent with the possibility that hippocampal recruitment is more closely related to accumulated multilingual experience than to aptitude per se. However, because this effect was not present in the full model including aptitude and the aptitude–multilingualism interaction, it should be interpreted cautiously.

Taken together, these results indicate that during the processing of intelligible native speech, both in the intact condition and in the high-level speech contrast, multilinguals show elevated activity outside what is typically considered the core perisylvian language network, also engaging regions associated with narrative comprehension and multimodal communication. These findings are consistent with the primary function of language in social interaction ^76^, particularly in multilinguals. Individuals who navigate multiple languages frequently may rely more strongly on alternative, multimodal cues and more extensively engage narrative-tracking regions to follow conversations in languages in which they have lower proficiency. Ultimately, the fact that this effect is detectable during native-language comprehension suggests that multilinguals may integrate these interactive and narrative systems with core language hubs more consistently than monolinguals.

As mentioned, degraded speech showed a different pattern of results, consistent with a greater top-down lexical and possibly syntactic decoding strategy in more multilingual individuals. Moreover, this pattern aligns with anecdotal reports during scanning from our most multilingual (i.e. polyglot) participants. Some of them reported enjoying the degraded speech blocks and attempting to decode them consciously, with a few claiming to have successfully identified words. Although decoding the snippets was impossible due to their unintelligibility (by design), the study highlights a strong tendency to project linguistic structure onto ambiguous auditory inputs. More multilingual participants showed increased activation within the left MTG, an area central to lexical processing ^77,78^, and within the IFG, in the right hemisphere in the full interaction model and bilaterally in the single-predictor model. The left IFG is a recognised hub involved in phonetic, lexical and morphosyntactic processing ^79–81^. The right IFG, in contrast, has been implicated in executive functions, particularly in inhibition ^82^ and in attention to relevant cues ^83^, and has been shown to be active during degraded speech processing in attentive listeners ^38^. It has also been identified as a central hub in language processing for bilinguals with experience in tonal languages ^84^. This pattern could reflect greater top-down decoding effort: because highly multilingual individuals possess broader linguistic repertoires, they may have a larger pool of potentially relevant lexical and structural candidates to actively navigate and select from when parsing an ambiguous acoustic signal. At the same time, higher multilingualism was accompanied by lower activity in left medial frontal regions, and hippocampal/parahippocampal regions (i.e. declarative memory hubs). Rather than indicating a simple increase or decrease in overall effort, this pattern may reflect a redistribution of processing resources: more multilingual individuals may engage regions involved in cue-based linguistic selection and decoding, including IFG and MTG, while relying less on broader conflict-monitoring and declarative-memory systems during acoustic challenge.

While this pattern of results warrants further investigation, it underscores the need for closer scrutiny of control conditions in language localiser paradigms. In typical localiser studies, these activations are routinely subtracted out, creating the risk of removing meaningful language-related activation that is tied directly to the participant’s linguistic phenotype and to the core scientific question. The issue may be particularly relevant in studies of multilinguals or high-aptitude participants, since both could be expected to engage more attentively with speech-like stimuli. In these cases, activation during degraded speech or other routinely used control conditions, such as nonwords, may be directly tied to decoding attempts and the engagement of linguistic processes. These results indicate that the adoption of a universal language localiser is a complex endeavour. Standardised, ready-to-use localisers are practical for clinical work, challenging populations, or general research purposes; however, targeted neurolinguistic investigations require experimental tasks and control baselines that are carefully tailored to the specific research questions and populations under study.

Finally, the lack of significant relationships between multilingual experience and activation in areas traditionally associated with bilingual and multilingual experience during intact speech processing and the high-level speech contrast processing was surprising. We observed no robust effects of multilingualism within classical language processing hubs or networks typically tied to language control and switching. This absence of an effect may be due to the characteristics of our sample, which consisted of mostly multilingual participants, with very few individuals falling towards the strictly monolingual end of the spectrum. This absence of consistent effects in these regions could also reflect the variability of real-world multilingual practices and profiles, where factors such as switching frequency, environmental context, and domain-general mechanisms supporting multilingual language use may leave a more variable neural signature.

### c. Aptitude is related to efficient recruitment of language hubs

Language aptitude exhibited a neural signature characterised primarily by lower activation in core language regions among individuals with higher aptitude during intact speech processing. Specifically, this lower recruitment was restricted to left-hemisphere structures, such as the IFG, MTG and angular regions. The left angular gyrus has been linked to aptitude for vocabulary learning ^85^, together with the left MTG ^86^, as well as to more efficient grammar learning ^87^. During the high-level speech contrast, this pattern of reduced activity extended to bilateral clusters, with a distinct left-hemisphere bias in frontal regions, alongside lower activation in primary auditory and superior temporal areas. Individual differences in the anatomy of auditory areas such as HG have already been shown to be linked to aptitude for foreign speech-sound perception ^88,89^ and production ^90^, as well as to measures of different aptitude subcomponents, including phonological, morphosyntactic, and rote-learning skills ^48,91,92^. Moreover, while previous functional neuroimaging research has largely investigated language aptitude subcomponents separately to identify the neural correlates of individual skills, our work leveraged a single compound score, which nevertheless proved sensitive to activation patterns in regions previously associated with different components of language aptitude, including phonological, morphosyntactic, and rote-learning abilities.

These results suggest that the neural efficiency previously attributed to polyglotism ^3,4^ may be more accurately associated with language aptitude than with multilingual experience, measured using either categorical or continuous approaches. While more multilingual speakers appear to engage a stronger non-language-specific network to manage their active linguistic repertoire, individuals with high aptitude appear to process linguistic information with fewer neural resources. This divergence could reflect a predisposition for language, whereby ease with language learning and use is marked by efficient cortical processing within dedicated language hubs during intelligible speech processing conditions. Our findings further support our hypothesis that prior investigations into high-level multilingualism, which are frequently are influenced by the Eurocentric view that sees monolingualism as the norm, as it is the case in many WEIRD countries, and treating multilingualism as the exception, possibly reserved for talented language learners, may bias our baseline understanding of how the brain manages multiple languages. A true understanding of the neurobiology of multilingualism should sample multilingual as well as monolingual context and/or take language aptitude into account, just as studies of language aptitude should consider multilingual experience. Given that the vast majority of bilingual and multilingual individuals globally acquire additional language due to migration or environmental necessity rather than elective choice or passion, accounting for these distinct dimensions yields a more ecologically valid framework that is more representative of the lived reality of multilinguals worldwide.

Lastly, the neural response to degraded speech highlights a fundamental divergence between more multilingual and higher aptitude participants. Whereas multilingual individuals appear to draw on their extended linguistic repertoire to engage in high-level, top-down decoding, reflected in higher activation of the IFG and MTG, high-aptitude learners may rely more strongly on the extraction of prosodic and auditory properties of the signal, as evidenced by modulation in the right STG and STS ^93,94^. This pattern may reflect a comparatively more bottom-up approach to parsing novel or ambiguous linguistic input.

### d. Operationalising multilingual experience: categorical grouping vs. continuous measures

Measuring multilingual experience using a continuous measure was better at capturing neural correlates related to variation in multilingual experience than categorising participants into the artificial categories of polyglots and controls. As shown in Figure 1, our participants – and we would argue multilinguals more generally – are better represented along a continuum than by an over-simplified dichotomous categorisation. Thus, it is not surprising that a continuous approach was better able to detect differences in neural processing during native speech comprehension. We also did not reproduce the findings from previous studies of polyglots ^3,4^, even when applying similar categorisation criteria and, in a second analysis, using the same subject-specific fROI approach. While the lack of convergence in the whole-brain analysis may partly reflect differences in statistical approach, namely whole-brain voxel-wise analysis versus subject-specific functional ROI analysis, the absence of significant results when using the previously adopted subject-specific fROI analysis likely reflects differences between the samples themselves. In the present work, we captured a broad, diverse spectrum of both aptitude and multilingualism, but with an under-representation of monolinguals. It is also worth noting that our continuous characterisation of language background accounted for any language known at any level. If we had instead restricted our criteria to advanced proficiency, 64 of our participants would have been labelled as knowing only one language, effectively placing them in what previous studies consider the monolingual or control baseline. This issue is not specific to studies of polyglotism, but may also arise in bi-and trilingualism research, where partial knowledge of additional languages is often not reported or quantified.

## 4. Conclusions

In summary, this study provides evidence for behavioural and neural dissociations between the effects of language aptitude and multilingual experience on neural recruitment during speech processing. While language aptitude was associated with a neural profile consistent with efficiency, marked by reduced activation within core perisylvian language hubs, multilingual experience was associated primarily with activation differences outside these traditional networks, engaging areas associated with narrative tracking, gesture-related processing, and multimodal communication. This suggests that a listener’s predisposition for language learning is reflected in core speech-processing hubs, whereas experience with multiple languages is associated with greater recruitment of regions supporting narrative tracking, gesture-related processing, and multimodal aspects of communication, rather than primarily altering core linguistic nodes.

Moreover, evaluating individual task conditions rather than relying solely on subtraction contrasts proved critical, suggesting that highly multilingual individuals may actively attempt to decode and extract meaning even from degraded, unintelligible auditory input.

Finally, the significant interaction detected between these two dimensions underscores that examining aptitude or multilingualism in isolation misses the complex neural effects that emerge only when language-learning predisposition and multilingual experience are modelled together and when sample characteristics, particularly regarding the nature of multilingual experience, are carefully contextualised. Ultimately, these findings demonstrate that a more complete understanding of multilingual neurobiology, and a true delineation of the neural signatures of innate predisposition and experience, requires an integrated approach that models these factors in tandem rather than in isolation.

## 5. Methods

### a. Participants

Data from 121 participants (mean age 24.9, *SD*=6.5, 82 females) were included in this study. This comprised the openly available NEBULA 101 dataset ^40^ and data from an additional 20 healthy polyglots. The participants spanned a wide range in terms of their multilingual language experience, as shown in Figure 1 and Table 2. Participants completed a series of behavioural tasks, questionnaires and neuroimaging measures described in depth and analysed in other works ^41,42^. For the purposes of the present work, we describe only relevant demographics, multilingualism and aptitude measures. All participants provided signed informed consent. The study received ethical approval from the Geneva Cantonal Ethical Commission (Protocol N. 2021–01004).

### b. Behavioural tasks and analysis

#### i. Multilingualism

Multilingual experience was measured through self-report using the Leap-Q questionnaire ^43^. Participants self-reported their proficiency (on a scale from 1 to 10) in speaking, writing and comprehension for each language they knew. For the continuous multilingualism score, self-reported speaking proficiency was used to calculate an entropy measure ^44^. Entropy measures indicate the degree of multilingualism by capturing both the number of languages spoken and the balance of proficiency across them. Such continuous measures offer a more ecologically valid alternative to categorical grouping, as they capture both the spread and level of language proficiency for each speaker ^45^, thereby better representing the graded nature of multilingual language use.

In addition to the continuous approach described above, we also divided participants into polyglots and controls, following the grouping criteria used in the two previous fMRI studies of polyglots ^3,4^. We ran additional analyses using a binary group comparison, with the aim of replicating this previously published work on polyglots. This resulted in 41 participants categorised as polyglots (mean age 28.8, SD=8.8, 26 females), with 14 qualifying as hyperpolyglots, and 80 categorised as controls (mean age 23.0, SD=3.6, 56 females). More information on categorisation and grouping outcomes is available in the Supplementary Materials (Table s1, Figure s1.)

#### ii. Language aptitude tasks

To assess language aptitude, we tested participants on measures of foreign speech sound perception and production, grammatical sensitivity, and rote learning ability. In-depth descriptions of each task are available in the dataset publication ^40^ and other previous works ^41,42^. Table 1 provides a quick overview of the tasks and single-task score calculations. For the brain-behaviour analysis, aptitude task scores were z-scored against the sample mean and summed to obtain one comprehensive aptitude index for each participant.

**Table 1.** Overview of the tasks and scores used to calculate the compound language aptitude score.

| Task name | Description | Score | Construct |
| --- | --- | --- | --- |
| Farsi uvular sound production | Imitation of foreign speech sounds | Percentage of correct responses (2 native raters) | Foreign speech sound production |
| Hindi dental-retroflex sound categorisation | Adaptive task measuring ability to discriminate the dental-retroflex plosive consonant foreign speech sound contrast | Accuracy weighted by difficulty | Foreign speech sound perception |
| ArtGram | Ability to infer morphosyntactic characteristics of an artificial language and translate sentences after presentation of examples and a dictionary | Accuracy | Grammatical sensitivity |
| MLAT 5 | Vocabulary learning in an unknown language (MLAT 5) | Accuracy | Rote learning |

**Table 2.** Descriptive statistics for behavioural variables of interest: multilingualism and language aptitude (*N* = 121).

| Measure | *M* | *SD* | *Min* | *Max* |
| --- | --- | --- | --- | --- |
| Multilingualism (speaking entropy) | 1.39 | 0.52 | 0.00 | 3.70 |
| Aptitude (z-scored) | 0.00 | 2.30 | -6.25 | 4.74 |
| Number of languages known | 6.27 | 5.71 | 1.00 | 50.00 |
| Number of languages known at advanced proficiency | 1.96 | 1.69 | 1.00 | 11.00 |

**Table 3.** Multiple linear regression: relationship between multilingualism and language aptitude.

| Predictor | $\beta$ | SE | t | p | 95% CI [LL, UL] |
| --- | --- | --- | --- | --- | --- |
| Intercept | 0.173 | 0.238 | 0.724 | .470 | [-0.300, 0.645] |
| Language Aptitude | 0.038 | 0.017 | 2.313 | .022 | [0.006, 0.071] |
| Sex (Binary) | -0.073 | 0.083 | -0.869 | .386 | [-0.238, 0.093] |
| Age | 0.050 | 0.007 | 6.819 | < .001 | [0.035, 0.064] |
| Education | -0.000 | 0.018 | -0.015 | .988 | [-0.036, 0.035] |

**Table 4.**
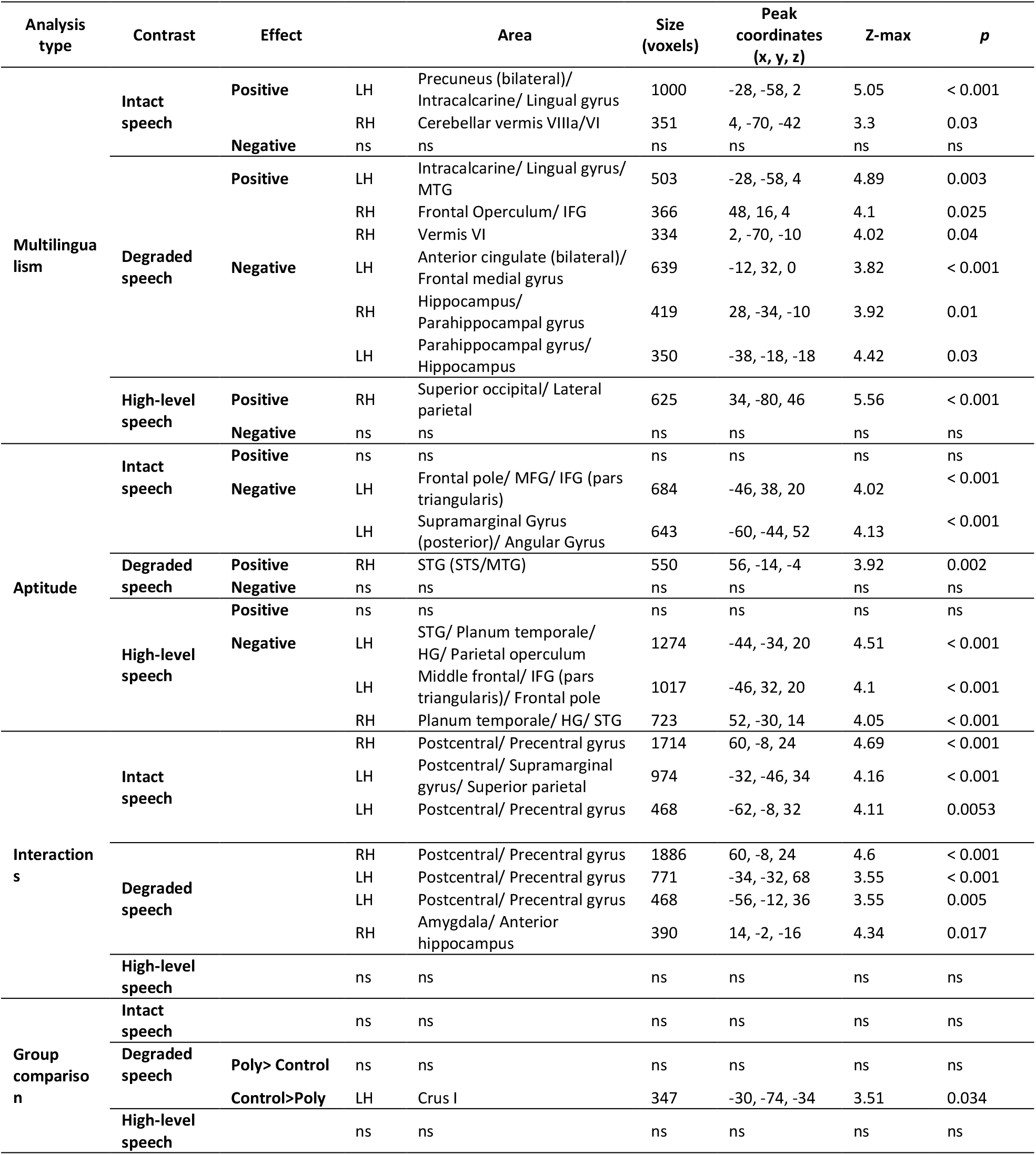
Brain activity results showing significant associations with multilingualism, language aptitude, their interaction, and categorical group membership across the functional contrasts. Regions are labelled using visual inspection and according to the most probable anatomical label based on the Harvard-Oxford cortical and subcortical atlases and the FNIRT cerebellar atlas. ns = non-significant. IFG = inferior frontal gyrus, MTG = middle temporal gyrus, STG = superior temporal gyrus, STS = superior temporal sulcus, HG = Heschl’s gyrus.

#### iii. Assessment of the dissociation between language aptitude and multilingualism

To evaluate the degree of association between our primary predictors, we conducted a partial correlation between language aptitude and multilingual entropy while controlling for age, sex, and education. We subsequently performed a multiple linear regression to determine the relationship between language aptitude and multilingual experience when accounting for these same demographic covariates.

### c. Speech processing in the brain

#### i. MRI measures

For this study, we made use of brain structural MRI (MPRAGE) and functional MRI (fMRI) data acquired during a language localiser (see Rampinini et al, 2025 for more details). The localiser was administered both in the participants’ first (L1) and second (L2) languages. In this work, we analysed only fMRI data from the L1 localiser condition. Images were acquired using a Siemens 3T Magnetom-Prisma scanner at Campus Biotech, Geneva, equipped with a 64-channel head coil. T1-weighted MPRAGE structural images were collected using 208 sagittal slices (TR = 2,300 ms, TE = 3.26 ms, voxel size = 1 × 1 × 1 mm). BOLD activation was measured using a whole-brain EPI sequence (acceleration factor = 1, FOV = 224 mm, voxel size: 2 × 2 × 2 mm, TR = 2,000 ms, TE = 32 ms, flip angle = 75 °, interleaved acquisition, 72 slices).

During the language localiser task, participants passively listened to intact and degraded speech passages from the story *Alice in Wonderland* while looking at a fixation cross on a white screen. The stimuli and task are openly available and described in previous works ^4,10^. We used the localiser stimuli and set up as provided in the openly available task ^10^ but we modified the task presentation in order to present both intact and degraded passages in the L1 (usually French, N=111) and the L2 to each participant. Degraded passages were unintelligible and resembled a poor radio transmission. They were obtained by low-pass filtering and combined with amplitude envelope-modulated noise, resulting in the removal of phonological and linguistic content (details of the degradation procedure described in Scott et al., 2017). In total, three runs were acquired, each including five baseline blocks consisting of 12 seconds of silent fixation and 16 task blocks of 18 seconds each, with four blocks for each language-by-condition combination. No response was required, and participants took short, self-timed breaks between blocks.

#### ii. Preprocessing of fMRI data

Preprocessing was carried out using FEAT (FMRI Expert Analysis Tool, version 6.0) within FSL 6.0 (FMRIB’s Software Library) using a standard preprocessing pipeline that included the following steps: removal of non-brain tissue using optiBET (Lutkenhoff et al. 2014), B0 unwarping using fieldmap images, non-linear registration to structural T1 images (BBR, FNIRT), and spatial smoothing using a Gaussian kernel of 5 mm FWHM. Data were motion-corrected using ICA-AROMA (Pruim et al. 2015), and no participant exceeded the absolute displacement threshold of 5 mm, which would have resulted in exclusion from the sample. We then removed nuisance regressors for white matter (WM), and cerebrospinal fluid (CSF), performed high-pass filtering at 90 s, and registered the preprocessed data to the MNI152 standard space (Grabner et al. 2006).

#### iii. First level modelling and brain-behaviour analysis

The BOLD response was modelled using a double-gamma haemodynamic response function. Three contrasts of interest were generated by calculating activation for: (i) intact blocks vs. baseline (intact speech), (ii) degraded blocks vs. baseline (degraded speech), and (iii) intact vs. degraded blocks. This latter subtraction contrast is often referred to in the literature as indexing high-level language processing, or simply as identifying the ‘language network’, because it is intended to isolate responses to intelligible linguistic input, particularly lexico-semantic and morphosyntactic processing, while removing lower-level auditory and modality-specific responses shared with the degraded condition. In the present study, we refer to this as the ‘high-level speech’ contrast. This terminology acknowledges that the contrast may reflect not only high-level language features but possibly also some activation related to phonological processing, which is largely absent in the degraded speech used as a control condition. To investigate how multilingualism, measured continuously, relates to neural activation, we performed a GLM analysis in FEAT FLAME (FMRI Expert Analysis Tool, version 6.0). We ran a full interaction model using multilingualism and aptitude scores and their interaction as predictors of interest, while age, sex, education and handedness were included as covariates. All scores were mean centred for interpretability. For completeness and to better interpret the stability of these effects, we also ran two single-predictor models, one including only multilingualism and one including only aptitude, controlling for the same covariates. Activation estimates for each of these three contrasts served as the dependent variables. To reproduce the group-level analysis approach used in previous studies of polyglot speech processing using a localiser approach, we also analysed activation differences based on categorical group membership. Specifically, we compared activation across the three contrasts between the polyglot group and a larger control group, using the same control variables (age, sex and education) as in the behavioural analysis (section 2.b.iii) with the addition of handedness, because of its potential relevance for hemispheric language organisation and functional activation patterns during speech processing. All results were thresholded using a cluster-forming threshold of Z > 2.3, with a cluster-corrected significance threshold of *p* =.05.

Finally, we performed a GSS ROI (fROI) analysis ^47^, an approach used in previous functional localiser studies of polyglots ^3,4^. This approach identifies the 10% most active voxels within a set of predefined ROIs for each participant (bilateral perisylvian regions; mask available at https://www.evlab.mit.edu/resources-all/download-parcels), allowing us to calculate individual effect sizes for each condition as well as the overall extent of activation. Although not the primary focus of the present study, we used these subject-specific metrics to assess whether previously reported group-level findings could be reproduced in our sample, again controlling for age, sex, education, and handedness.

## 6. CRediT authorship contribution statement

I.B: Conceptualization, Data curation, Formal Analysis, Investigation, Methodology, Project administration, Visualization, Writing – original draft, and Writing – review & editing. O.K.: Conceptualization, Data curation, Formal Analysis, Visualization, Writing – review & editing. A.R.: Data curation, Project administration, Writing – review & editing. R.B.: Conceptualization, Funding acquisition, Writing – review & editing. N.G.: Conceptualization, Funding acquisition, Writing – review & editing.

## 7. Acknowledgments and Funding

We would like to thank Sophie Slaats for her insightful and valuable discussions concerning our results. We would like to thank Michael Erard, Richard Simcott, Alexandre Teiga, Susan Fitzgerald, Lane Greene, Hugues Pluvinage and Swiss Radiotelevision for their help in the recruitment of participants. We also thank Neiloufar Family and Sevil Maghsadhagh for rating our foreign speech sound imitation task. This work was supported by the Swiss National Science Foundation [Grant #100014_182381], and by the NCCR Evolving Language, Swiss National Science Foundation [Agreement #51NF40_180888].

## 8. Data and code availability

The majority of the raw data used in this publication are openly available in the NEBULA101 dataset ^40^. All analysis scripts, behavioural metrics, and data required to reproduce the behavioural and neural interaction visualizations are available on GitHub (10.5281/zenodo.21489012). Additionally, FSL group-level design files and extracted mean activity for statistically significant brain clusters are included in the same repository.

