## Supplementary Materials for "Aptitude, not polyglotism, is associated with efficient activation in core language areas"

### Group comparison

*Creation and characterisation of the group-analysis sample, with additional group-comparison results*

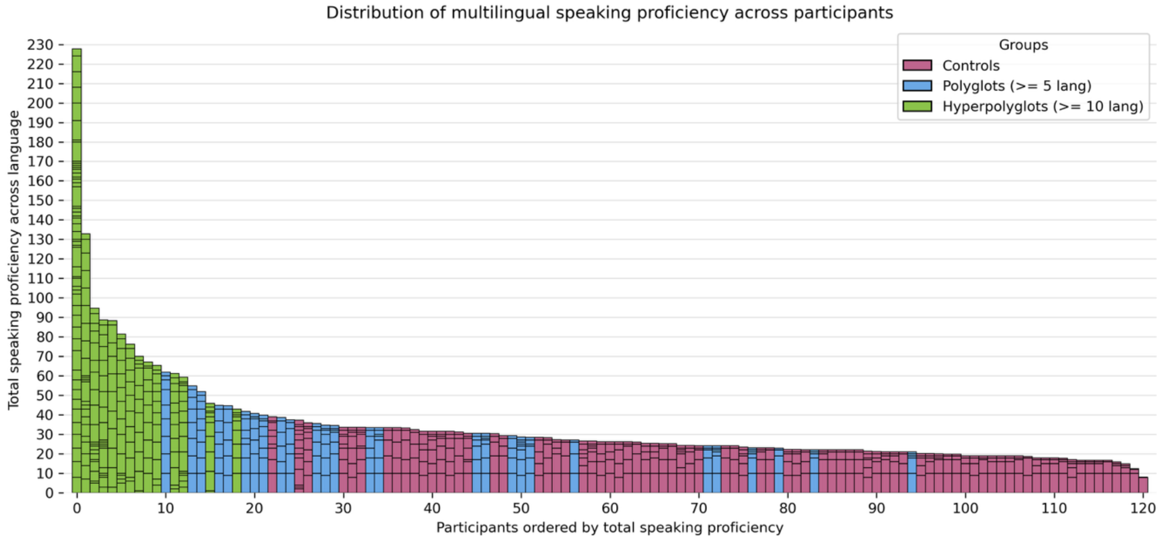


**Figure s1. Multilingual experience in the sample, colour-coded by group category.**

Each bar represents one participant's language experience; stacked segments within each bar represent self-reported speaking proficiency in individual languages. Total bar height therefore reflects cumulative speaking proficiency across languages, whereas the number of segments reflects the number of languages reported. Participants with the most extensive multilingual profiles, including hyperpolyglots, are shown on the left, whereas participants with less extensive multilingual experience, including monolinguals, are shown on the right.

| Variable | Polyglots (n=41) | Controls (n=80) |
|---|---|---|
| **Age (years)** | | |
| *M (SD)* | 28.78 (8.80) | 22.99 (3.56) |
| **Sex** | | |
| Female | 26 (63.4%) | 56 (70.0%) |
| Male | 15 (36.6%) | 24 (30.0%) |
| **Number of Languages** | | |
| *M* | 10.02 | 4.35 |
| *Mdn* | 7.00 | — |
| Range | 5.00–50.00 | 1.00–12.00 |
| **Hyperpolyglots (≥10 languages)** | 14 (34.1%) | — |

**Table s1. Demographic and language background characteristics of polyglots and controls**

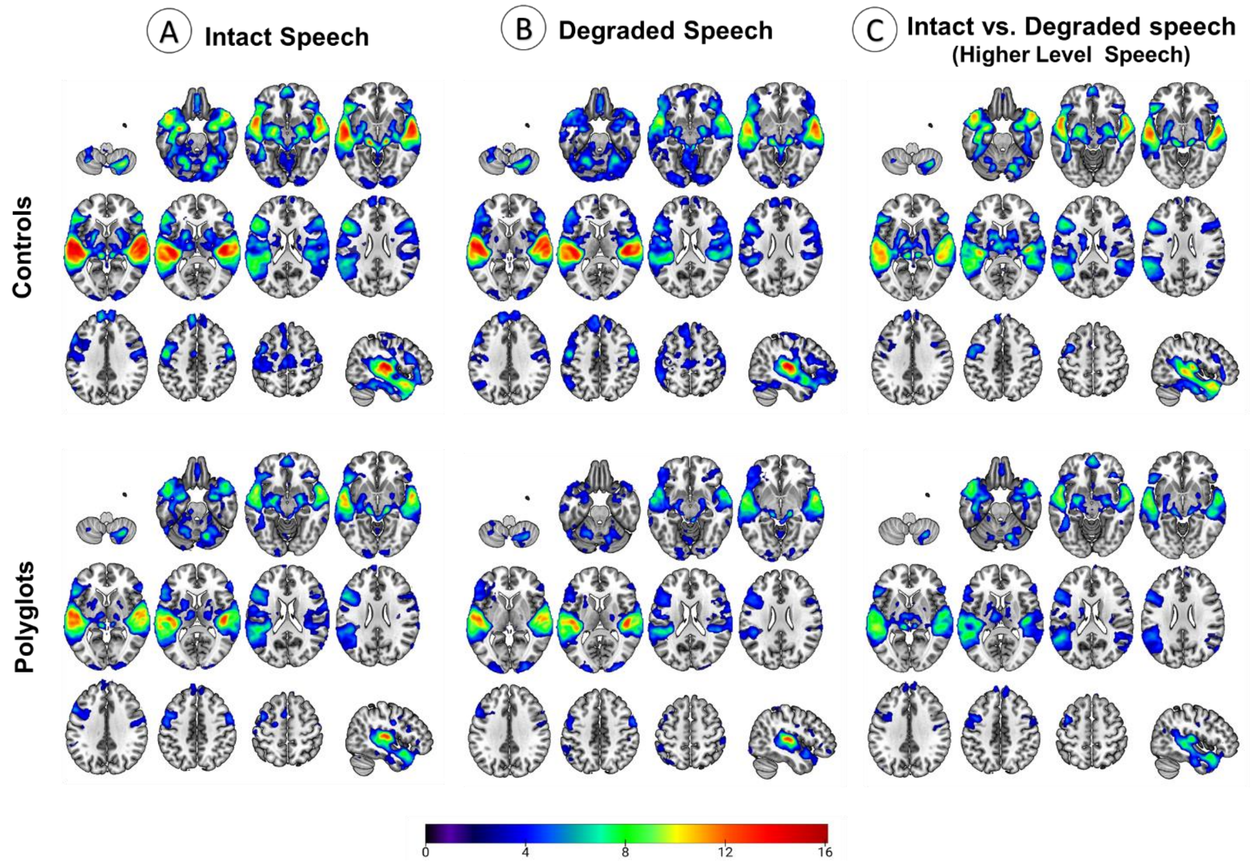


**Figure s2. Group activation in controls and polyglots across conditions**.

The colour bars indicate voxel-wise Z-score values for significant neural activation. Statistical images were thresholded using a cluster-forming threshold of Z > 2.3 and a cluster-corrected significance threshold of p = 0.05. Lateral view in all panels shows the left hemisphere.

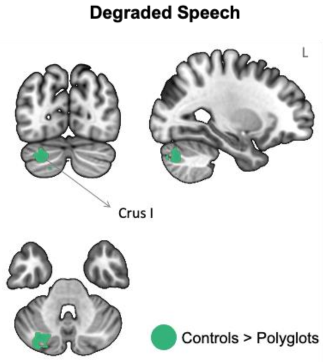


**Figure s3. Significant group difference between controls and polyglots during degraded speech processing.**

The cluster reflects greater activation in controls than polyglots. Statistical images were thresholded using a cluster-forming threshold of Z > 2.3 and a cluster-corrected significance threshold of p = 0.05.

**Single-predictor model results**

| Analysis type | Contrast | Effect | | Area | Size (voxels) | Peak coordinates (x, y, z) | Z-max | p value |
|---|---|---|---|---|---|---|---|---|
| **Multilingualism** | **Intact speech** | **Positive** | LH | Precuneus | 490 | 0, -82, 40 | 3.64 | 0.00385 |
| | | | LH | Intracalcarine/ Lingual gyrus | 404 | -28, -58, 4 | 4.92 | 0.0134 |
| | | | RH | Cerebellar vermis VIIIa/b | 339 | 6, -66, -40 | 3.39 | 0.0364 |
| | | **Negative** | ns | ns | ns | Ns | ns | ns |
| | **Degraded speech** | **Positive** | LH | Intracalcarine | 524 | -28, -58, 4 | 4.92 | 0.00251 |
| | | | RH | IFG (frontal operculum) | 416 | 48, 16, 4 | 4.18 | 0.0117 |
| | | | RH | Vermis VI | 366 | 2, -70, -10 | 4.11 | 0.0248 |
| | | | LH | IFG (pars opercularis) | 343 | -58, 14, 2 | 3.86 | 0.0354 |
| | | **Negative** | LH | ACC/ white matter | 749 | -12, 32, 2 | 3.82 | 0.000137 |
| | | | RH | Parahippocampal gyrus/ Hippocampus | 387 | 28, -34,12 | 3.64 | 0.018 |
| | | | LH | Parahippocampal gyrus/ Hippocampus | 342 | -18, -32, -10 | 4.27 | 0.036 |
| | **High-level speech** | **Positive** | RH | Superior lateral occipital cortex/ Angular gyrus | 661 | 40, -78, 44 | 5.78 | 0.000528 |
| | | | RH | Hippocampus/ Parahippocampal gyrus/ Inferior temporal | 380 | 40, -20, -18 | 4.27 | 0.0236 |
| | | **Negative** | ns | ns | ns | Ns | ns | ns |
| **Aptitude** | **Intact speech** | **Positive** | ns | ns | ns | Ns | ns | ns |
| | | **Negative** | LH | Frontal pole/ Middle frontal gyrus | 591 | -46, 38, 20 | 3.01 | <0.001 |
| | | | LH | Supramarginal gyrus/ Angular gyrus | 643 | -52, -46, 52 | 3.82 | 0.005 |
| | **Degraded speech** | **Positive** | RH | STG | 678 | 56, -16, -2 | 3.91 | <0.001 |
| | | **Negative** | ns | ns | ns | Ns | ns | ns |
| | **High-level speech** | **Positive** | LH<br>RH | Superior parietal lobule<br>Precuneus - | 494 | -26, -42, 52 | 3.41 | 0.0047 |
| | | **Negative** | LH | Posterior and anterior STG, planum temporale/ Planum polare/ HG | 879 | -62, -8, 4 | 4.4 | <0.001 |
| | | | LH | IFG (pars triangularis)/ Middle frontal gyrus | 708 | -46, 32, 20 | 4.17 | <0.001 |
| | | | RH | Posterior and anterior STG/ Planum temporale/ HG | 561 | 52, -30, 14 | 2.72 | 0.002 |
| **Group comparison** | **Intact speech** | | ns | ns | ns | Ns | ns | ns |
| | **Degraded speech** | **Poly > Control** | ns | ns | ns | Ns | ns | ns |
| | | **Control > Poly** | LH | Crus I | 347 | -30, -74, -34 | 3.51 | 0.0336 |
| | **High-level speech** | | ns | ns | ns | ns | ns | ns |

**Table s2. Significant clusters from the single-predictor models and categorical group comparison.** Note that one cluster (Aptitude/High-level speech/Positive) spans both hemispheres. Brain activity results showing significant associations with multilingualism and language aptitude, as well as significant polyglot–control group differences. Regions are labelled according to the most probable anatomical label based on the Harvard-Oxford cortical and subcortical atlases and the FNIRT cerebellar atlas. ns = non-significant.

**Group comparison results from the subject-specific functional ROI (fROI) analysis**

| | Extent | | Effect Size | | | | | |
|---|---|---|---|---|---|---|---|---|
| | | | Intact | | Degraded | | Linguistic | |
| ROI | *t* | *p* | *t* | *p* | *t* | *p* | *t* | *p* |
| **AngG-LH** | 0.72 | 0.48 | -0.89 | 0.37 | -0.69 | 0.49 | -0.34 | 0.73 |
| **AngG-RH** | 1.48 | 0.14 | -0.93 | 0.35 | -1.47 | 0.15 | 0.61 | 0.54 |
| **IFG-LH** | 0.54 | 0.59 | -0.44 | 0.66 | -1.58 | 0.12 | 0.73 | 0.47 |
| **IFG-RH** | 0.28 | 0.78 | -0.97 | 0.33 | -1.16 | 0.25 | 0.11 | 0.91 |
| **IFGorb-LH** | 0.80 | 0.43 | -0.90 | 0.37 | -1.32 | 0.19 | 0.30 | 0.76 |
| **IFGorb-RH** | 0.77 | 0.44 | -1.23 | 0.22 | -1.06 | 0.29 | -0.35 | 0.72 |
| **MFG-LH** | 0.96 | 0.34 | 0.45 | 0.65 | -0.62 | 0.53 | 1.01 | 0.32 |
| **MFG-RH** | 0.57 | 0.57 | -0.30 | 0.76 | -0.83 | 0.41 | 0.49 | 0.63 |
| **antTemp-LH** | 1.19 | 0.24 | -0.55 | 0.59 | -1.68 | 0.10 | 0.83 | 0.41 |
| **antTemp-RH** | 0.00 | 1.00 | -1.60 | 0.11 | -2.02 | 0.045 | -0.13 | 0.90 |
| **postTemp-LH** | 1.16 | 0.25 | 0.39 | 0.70 | -0.61 | 0.54 | 0.99 | 0.32 |
| **postTemp-RH** | 0.01 | 0.99 | -1.04 | 0.30 | -1.20 | 0.23 | -0.28 | 0.78 |

**Table s3. Subject-specific fROI comparison between polyglots and controls.**

The table reports group differences in overall activation extent and condition-specific effect sizes for intact speech, degraded speech, and the linguistic contrast.

**Significant interaction effects**

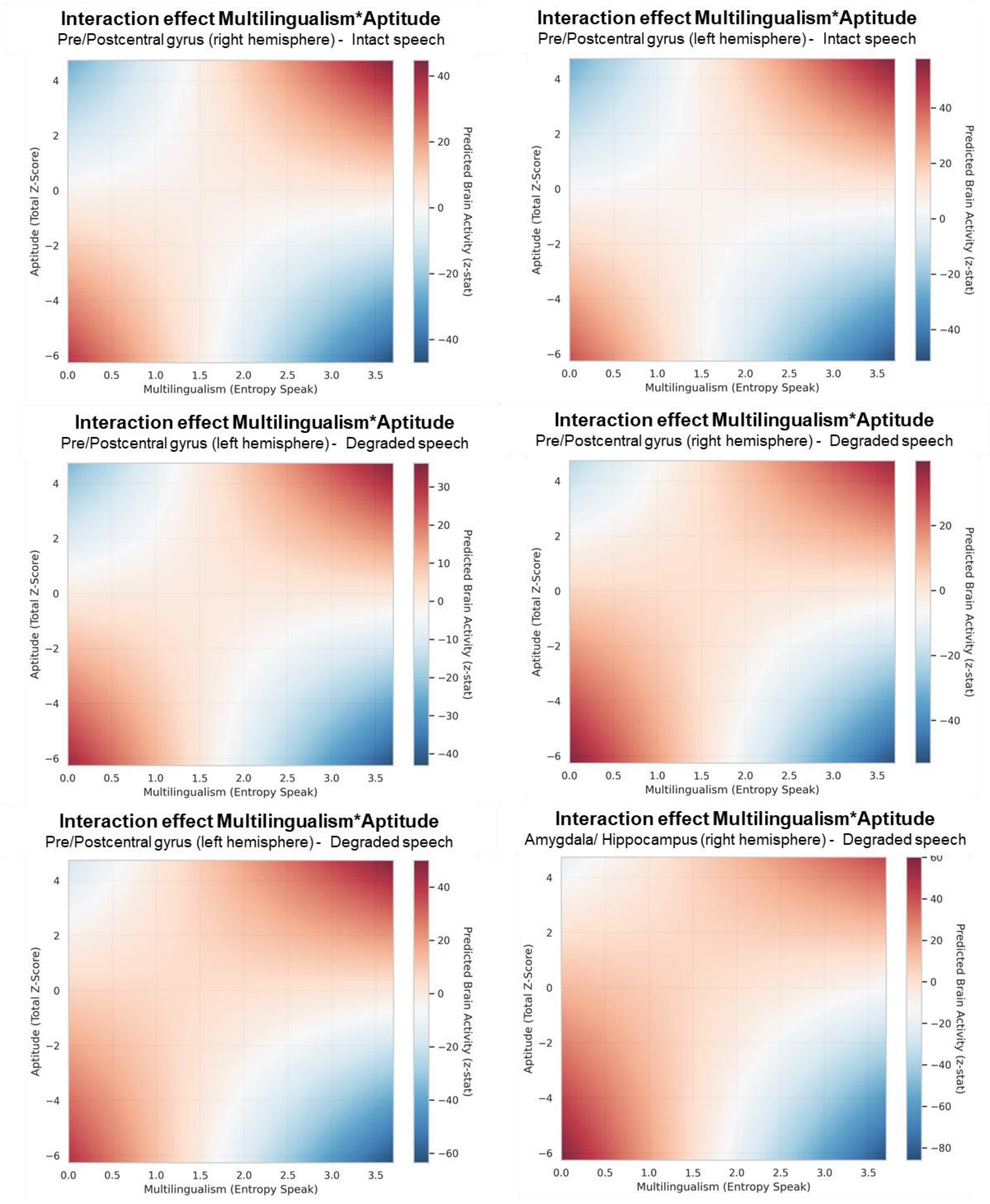


**Figure s4. Interaction between multilingual experience and language aptitude: post hoc decomposition of the interaction effect.**

Predicted neural activation within each ROI (average activation) is plotted as a function of continuous multilingual entropy and aptitude score.
